# Folliculin variants linked to Birt-Hogg-Dubé syndrome are targeted for proteasomal degradation

**DOI:** 10.1101/2020.03.30.015248

**Authors:** Lene Clausen, Amelie Stein, Martin Grønbæk-Thygesen, Lasse Nygaard, Cecilie L. Søltoft, Sofie V. Nielsen, Michael Lisby, Tommer Ravid, Kresten Lindorff-Larsen, Rasmus Hartmann-Petersen

## Abstract

Germline mutations in the folliculin (*FLCN*) tumor suppressor gene are linked to Birt-Hogg-Dubé (BHD) syndrome, a dominantly inherited genetic disease characterized by predisposition to fibrofolliculomas, lung cysts, and renal cancer. Most BHD-linked *FLCN* variants include large deletions and splice site aberrations predicted to cause loss of function. The mechanisms by which missense variants and short in-frame deletions in *FLCN* trigger disease are unknown. Here, we present computational and experimental studies showing that the majority of such disease-causing *FLCN* variants cause loss of function due to proteasomal degradation of the encoded FLCN protein, rather than directly ablating FLCN function. Accordingly, several different single-site FLCN variants are present at strongly reduced levels in cells. In line with our finding that FLCN variants are protein quality control targets, several are also highly insoluble and fail to associate with the FLCN-binding partners FNIP1 and FNIP2. The lack of FLCN binding leads to rapid proteasomal degradation of FNIP1 and FNIP2. Half of the tested FLCN variants are mislocalized in cells, and one variant (ΔE510) forms perinuclear protein aggregates. A yeast-based screen revealed that the deubiquitylating enzyme Ubp15/USP7 and molecular chaperones regulate the turnover of the FLCN variants. Lowering the temperature to 29 °C led to a stabilization of two FLCN missense proteins, and for one variant (R362C), FLCN function was re-established at low temperature. In conclusion, we propose that most BHD-linked *FLCN* missense variants and small in-frame deletions operate by causing misfolding and degradation of the FLCN protein, and that stabilization of certain disease-linked variants may hold therapeutic potential.

## Introduction

In order to function, most proteins require some conformational flexibility and are therefore not completely rigid in their native environment. In addition, since nature selects for function most proteins are not overly stable, and stress conditions or mutations may cause proteins to lose their native conformation and misfold. Because misfolded proteins can be toxic and form insoluble aggregates, they must be quickly eliminated from the cell via the so-called protein quality control (PQC) network (Hartl et al., 2011; Jones and Gardner, 2016; Kriegenburg et al., 2012). Typically, PQC depends on molecular chaperones to associate with the misfolded proteins, and refold them or guide them for degradation via the ubiquitin-proteasome system (UPS) or autophagy (Amm et al., 2014; Esser et al., 2004; Kevei et al., 2017; Shiber and Ravid, 2014; Sontag et al., 2014). Despite a number of recent efforts (Geffen et al., 2016; Maurer et al., 2016), it is not completely understood how cells discern misfolded from native proteins but it is likely to involve recognition of exposed hydrophobic regions that are normally buried in the native protein.

Recently, we have shown for several disease-related proteins that the degree of protein destabilization correlates with the turnover rate (Abildgaard et al., 2019; Nielsen et al., 2017; Scheller et al., 2019), and a structural destabilization of as little as 3 kcal/mol is enough to trigger degradation. This is in agreement with genetic studies in yeast that have shown that the PQC system operates by following a better-safe-than-sorry principle and is thus highly diligent and prone to target proteins that are only slightly structurally perturbed but still functional (Gardner et al., 2005; Kampmeyer et al., 2017a; Kriegenburg et al., 2014). Likewise, seminal studies in human cells have shown that some cystic fibrosis patients carry mutations in the *CFTR* gene that result in a protein that retains biochemical function. This protein variant fails to conduct its function not because it is intrinsically inactive, but because it is targeted by the PQC system for proteasomal degradation, which in turn leads to an insufficient amount of protein and ultimately disease (Ahner et al., 2007; Meacham et al., 2001).

Here, we show that a similar PQC mechanism is responsible for targeting certain variants of the folliculin protein (FLCN) linked to Birt-Hogg-Dubé (BHD) syndrome (OMIM: 607273) and that, similar to results for multiple other proteins, structural calculations predict these effects (Abildgaard et al., 2019; Arora et al., 2017; Beaver et al., 2019; Casadio et al., 2011; Caswell et al., 2019; Iqbal et al., 2019; Jepsen et al., 2019; Kumar et al., 2017; McCafferty and Sergeev, 2016; Nielsen et al., 2017; Pal and Moult, 2015; Scheller et al., 2019; Tang et al., 2019; Tiberti et al., 2019; Wagih et al., 2018). The autosomal dominantly inherited BHD syndrome is caused by mutations in the *FLCN* gene (Nickerson et al., 2002; Schmidt and Linehan, 2018; Zbar et al., 2002). BHD syndrome predisposes patients to develop hair follicle hamartomas (fibrofolliculomas), lung cysts, and renal neoplasia with an age-dependent and variable penetrance (Birt et al., 1977; Kovacs et al., 2008; Menko et al., 2009; Toro et al., 1999). A locus for BHD syndrome was mapped to chromosome 17p11.2 by linkage analysis in BHD families, and subsequently germline mutations in the *FLCN* gene were identified (Khoo et al., 2001; Nickerson et al., 2002; Schmidt et al., 2001). Further analysis of tumors from patients with BHD syndrome revealed somatic *FLCN* second hit inactivating mutations in the wildtype allele (Vocke et al., 2005; Yang et al., 2008), in line with FLCN having a tumor suppressor function. The tumor suppressor activity of FLCN is also supported by data from BHD animal models (Lingaas et al., 2003; Okimoto et al., 2004; Togashi et al., 2006).

The widely expressed *FLCN* gene encodes the 64 kDa cytosolic and nuclear FLCN protein, which does not display high sequence similarity to any other human protein but is highly conserved across species. Functional studies have linked FLCN to diverse metabolic pathways and cellular processes (Schmidt and Linehan, 2018), including the mTOR signalling pathway (Baba et al., 2008), regulation of PGC1α and mitochondrial biogenesis (Klomp et al., 2010), TFE3/TFEB transcriptional regulation (Hong et al., 2010), RhoA activation, cell adhesion (Nahorski et al., 2012), and amino acid-dependent activation of mTORC1 through Rag GTPases (Bar-Peled and Sabatini, 2014; Tsun et al., 2013).

A high-resolution crystal structure of the C-terminal domain of FLCN has been solved (Nookala et al., 2012) and shows a remarkable structural similarity to the DENN domain in the DENN1B protein (Wu et al., 2011). The DENN domain proteins have been found to function as guanine exchange factors (GEFs) for Rab-type GTPases and thus activate the vesicle transport function of these small GTPases (Yoshimura et al., 2010). Recently, a full-length structure of FLCN was obtained by cryo-EM (Shen et al., 2019), showing that the N-terminal region of FLCN contains a Longin domain which forms contacts with the Longin domain in its binding partner folliculin-interacting protein (FNIP1), while the C-terminal DENN domain interacts with the DENNc domain in FNIP1. In turn, FNIP1 mediates interaction to RagA, RagC and Ragulator to control mTORC activation (Shen et al., 2019). In addition to FNIP1, FLCN also associates with the related protein FNIP2 and the AMP-activated protein kinase AMPK (Baba et al., 2006; Hasumi et al., 2008; Takagi et al., 2008), which in cells serves as an energy sensor and regulator of the mTOR signalling pathway. The FNIP1 and FNIP2 proteins display sequence similarity (49% identity) and similar expression patterns (Hasumi et al., 2008), suggesting that they may overlap in function. The link between FLCN function and the FNIP1/2 proteins is also supported by genetic data. Thus, mice inactivated in both *Fnip1* and *Fnip2* develop cystic kidneys similar to the phenotype observed in kidney-targeted *Flcn* knock-out mice (Baba et al., 2008; Chen et al., 2008; Hasumi et al., 2015).

The vast majority of BHD-linked *FLCN* mutations include large deletions and splice site aberrations predicted to cause loss of function (Schmidt and Linehan, 2018). On the other hand, the consequences of the more rare BHD-linked *FLCN* missense variants and short in-frame deletions are not well characterized, although some are likely to destabilize the FLCN protein (Nahorski et al., 2011).

Here, we show that disease-causing *FLCN* missense mutations, rather than directly ablating FLCN protein function, instead render the FLCN proteins to become PQC targets. Rapid proteasomal degradation of the FLCN variants results in strongly reduced steady-state FLCN levels in cells. Several of the variants are also insoluble and fail to associate with the FLCN binding partners FNIP1 and FNIP2. This lack of FLCN binding in turn triggers proteasomal degradation of FNIP1 and FNIP2. Some of the FLCN variants are mislocalized in cells and one variant (ΔE510) forms perinuclear protein aggregates. The USP7 deubiquitylating enzyme (DUB) regulates FLCN turnover. In agreement, with the disease-linked FLCN variants being structurally destabilized, lowering of the temperature leads to a stabilization of the missense proteins, and for one variant function was reestablished at low temperature. In conclusion, we propose that BHD-linked missense mutations operate by causing misfolding and degradation of the FLCN protein, and structural stabilization of certain disease-linked variants may hold therapeutic potential.

## Results

### Disease-associated FLCN variants display reduced steady-state levels due to proteasomal degradation

Previous studies indicate that BHD-linked FLCN missense protein variants are destabilized (Nahorski et al., 2011), which, in turn, suggest that they might be subject to PQC-dependent clearance. To test this hypothesis, we first analyzed the steady-state level of wild-type FLCN and eight different FLCN variants that have been found in BHD patients (Table 1). The proteins were produced untagged from the pIRES2-GFP expression vector in transiently transfected U2OS cells. In agreement with the previous report (Nahorski et al., 2011), we found that several of the FLCN variants were present at strongly reduced steady-state levels (Fig. 1AB). The reduction in the FLCN protein levels was not caused by changes in transfection efficiency or in transcription, since the level of GFP, produced from the internal ribosomal entry site (IRES) in the same mRNA as FLCN, was unchanged (Fig. 1A). The most severe reduction in steady-state level was observed for the ΔE510 FLCN variant, while the levels of the V400I and K508R variants were hardly affected, compared to the wild-type FLCN control (Fig. 1AB). Indeed, although both V400I and K508R have been found in BHD patients, these occurred together with other putative pathogenic variants and their link to the disease is unclear (Table 1).

**Table 1.**

| *Stability, localization and function of FLCN variants* |  |  |  |  |  |  |  |  |  |
| --- | --- | --- | --- | --- | --- | --- | --- | --- | --- |
| FLCN variant | Protein level (%) | Half-life (hours) | Subcellular localization | Solubility | Stabilizes FNIP1/2 | Binds to FNIP1/2 | Stabilized at 29 °C | ClinVar category | Rosetta $\Delta\Delta G$ (kcal/mol) |
| WT | 100 | >8 | Cyt & Nuc | Normal | Yes | Yes | - | Benign | 0 |
| $\Delta F157$ | ~33 | ~3 | Cyt | Reduced | No | Poorly | No | Pathogenic | - |
| R239C | ~27 | ~3 | Cyt & Nuc | Very low | No | Yes | Yes | Conflicting | - |
| H255P | ~29 | ~3 | Cyt | Reduced | No | Poorly | Partially | VUS | - |
| R362C | ~59 | ~8 | Cyt & Nuc | Reduced | Partially | Yes | Yes | VUS | 2.9 |
| V400I | ~80 | >8 | Cyt & Nuc | Normal | Yes | Yes | - | Conflicting | 0.7 |
| $\Delta K508$ | ~31 | ~4 | Cyt | Very low | No | Poorly | No | Pathogenic | 5.0 |
| K508R | ~99 | >8 | Cyt & Nuc | Normal | Yes | Yes | - | Like. benign | 0.02 |
| $\Delta E510$ | ~23 | ~6 | Aggregates | Very low | No | Poorly | No | Pathogenic | 4.0 |

**Fig. 1.**
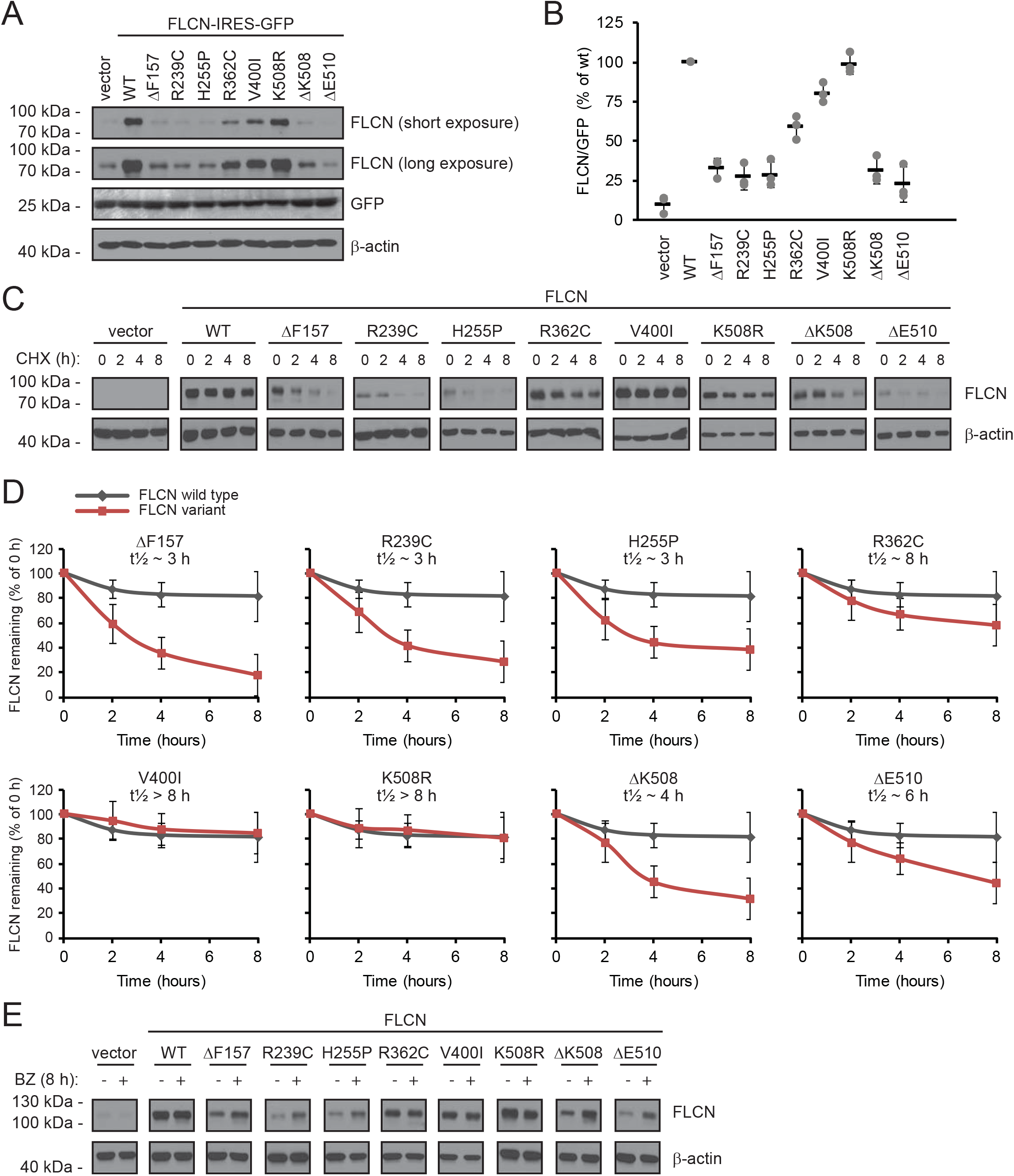
BHD-linked FLCN variants are rapidly degraded by the proteasome. (A) The protein level of wild-type (WT) FLCN and the FLCN variants in U2OS cells, was determined by SDS-PAGE and Western blotting with antibodies to FLCN. GFP and β-actin served as controls for equal transfection efficiency and loading, respectively. Endogenous FLCN was only visible upon longer exposures. (B) Quantification of blots as shown in (A) by densitometry normalized to the GFP level. The level of wild-type FLCN was set to 100%. The error bars show the standard deviation (n = 3). (C) The degradation of the FLCN variants was followed in cultures treated with the translation inhibitor cycloheximide (CHX) by SDS-PAGE and Western blotting with antibodies to FLCN. β-actin served as a loading control. (D) Quantification of blots as shown in (C) by densitometry normalized to the level at 0 hours. The error bars show the standard deviation. (n = 3). (E) The steadystate levels of the FLCN variants were compared in cultures incubated with or without the proteasome inhibitor, bortezomib (BZ), for 8 hours. β-actin served as a loading control.

To test directly if the reduced protein levels were caused by degradation, we followed the turnover of the FLCN variants in cultures treated with the translation inhibitor, cycloheximide (CHX). The wild type, V400I and K508R variants were stable, while the other variants were degraded (Fig. 1C). By densitometry of Western blots, we estimate the half-life of wild-type FLCN to be ≫8 hours, while the short-lived variants displayed half-lives between 3 and 8 hours (Fig. 1D).

Most intracellular proteins are degraded via the proteasome or via the autophagy system. To analyze whether the observed FLCN turnover was caused by proteasomal degradation or via autophagy, we analyzed the level of the FLCN variants in cultures treated with the proteasome inhibitor bortezomib (BZ) or chloroquine (CQ), which inhibits autophagy. In case of wild-type FLCN and the stable variants, V400I and K508R, there was no effect of BZ (Fig. 1E), since these variants are not rapidly degraded. However, for all other variants the levels were increased in response to proteasome inhibition (Fig. 1E), suggesting that these variants are proteasome targets. We did not observe stabilization in the presence of CQ for any of the variants (Fig. S1).

### In silico analyses suggest that pathogenic FLCN variants are structurally destabilized

We performed biophysical calculations to estimate the effects of the different FLCN variants on protein stability change (ΔΔG). The ΔΔG values quantify the loss or gain of favourable or destabilizing molecular interactions in the protein conformation induced by the missense variant or deletion, with high ΔΔG values indicating greater destabilization and thus more likely recognition by the PQC system. The calculations are structure-based and were only carried out for the C-terminal domain of FLCN, for which high-resolution structural details are available (residues 341 to 566, PDB 3V42, see also Fig. 2A) (Nookala et al., 2012).

**Fig. 2.**
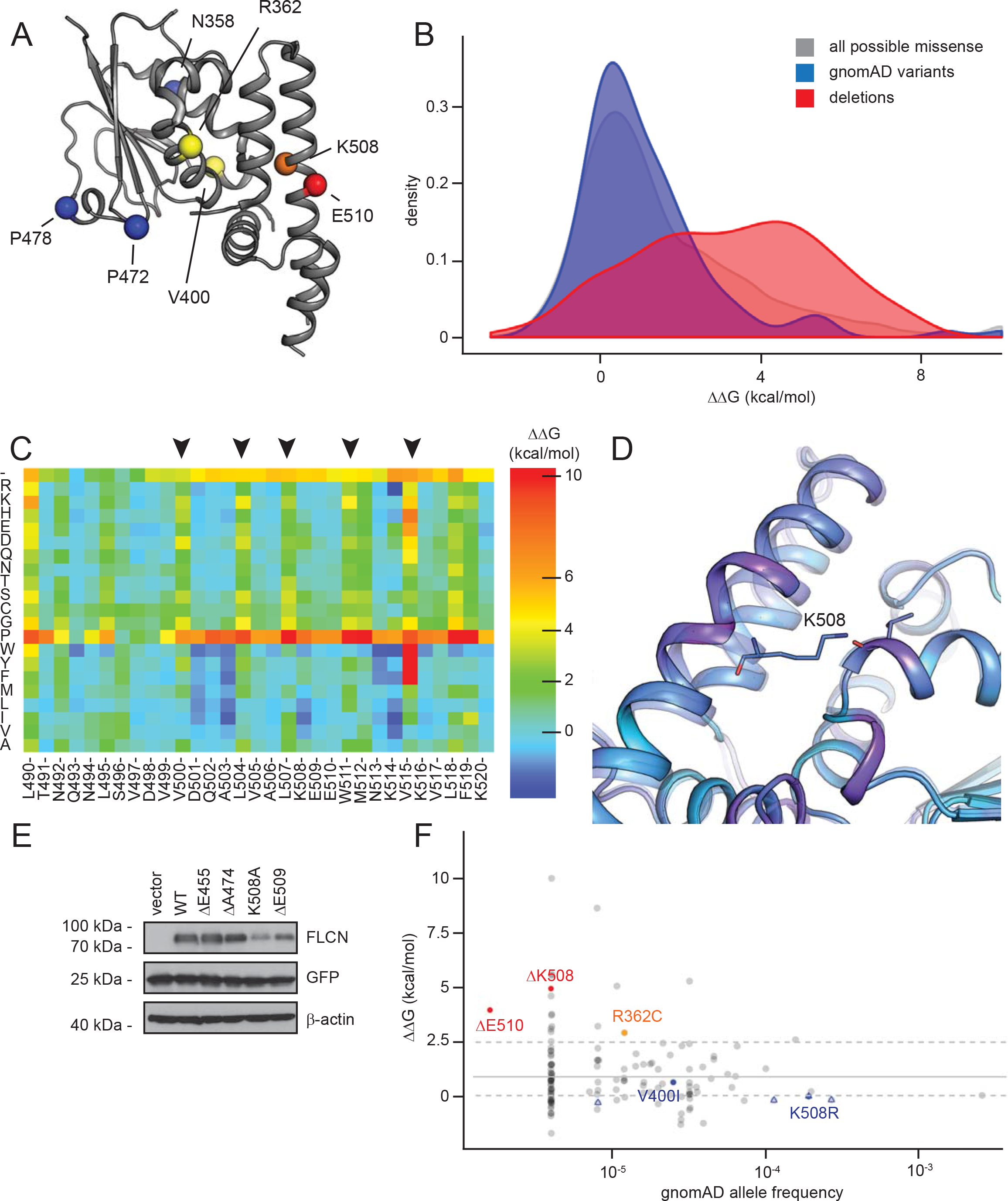
Structural consequences of the FLCN in-frame deletions. (A) Overview of the C-terminal domain of the FLCN crystal structure (PDB: 3V42) used for the structural calculations. Positions experimentally assayed in this work are indicated with spheres (red, deletions; yellow, missense variants; orange, position K508 with both missense variant and in-frame deletion; blue, benign ClinVar entries). (B) Distribution of ΔΔGs for all possible individual missense variants (gray), the subset of missense variants observed in gnomAD, and all possible individual deletions (red) in the C-terminal domain. For calculation details see Methods. (C) Heat map for the segment from position 490-520 visualizing both missense and deletion (-, top row) ΔΔGs. A large part of this sequence forms an alpha helix, and positions that point into the domain core (arrowheads) are more constrained than those pointing towards the solvent. (D) Structural model for the deletion of K508. Deletion model in solid tracing, wild-type transparent. The helix is strained (magenta) and the hydrogen bond which K508 forms with the backbone of T393 is lost. (E) The steady-state levels of the indicated FLCN variants were compared by blotting. β-actin served as a loading control. These additional variants were tested based on stability calculations. K508A leads to loss of hydrophobic interaction and the side-chain-backbone hydrogen bond described in (D) and is thus predicted to be less stable. ΔE445 and ΔA474 are in linker regions and predicted to not substantially destabilize the structure. ΔE509 was predicted to be as destabilizing as its neighbors ΔK508 and ΔE510, however, experimentally it is only mildly destabilized. This may be because its side chain points into solvent, whereas the side chains of K508 and E510 make important contacts with other parts of the C-terminal domain of FLCN. (F) “Fishtail plot” illustrating the gnomAD allele frequency of missense and deletion variants vs. the Rosetta ΔΔG. Empty blue triangles represent benign ClinVar entries. Common and benign variants have near-neutral stability, while several pathogenic variants are predicted to lose stability.

For missense variants, it is well-established that many have neutral or slightly destabilizing effects, and only a fraction of the possible single amino acid changes are severely detrimental (Matreyek et al., 2018; Roscoe et al., 2013; Stein et al., 2019). In contrast, there is far less data on protein stability for in-frame deletions, though existing studies on certain model proteins show that a number of both insertions and deletions are functional (Arpino et al., 2014; Banerjee et al., 2019; Gonzalez et al., 2019; Jones, 2005). All three deletions in our initial dataset, however, are severely destabilised (Fig. 1). Thus, we aimed to identify putative non-deleterious deletion variants and validate those experimentally.

We calculated ΔΔGs for missense variants with Rosetta (Alford et al., 2017; Park et al., 2016; Scheller et al., 2019) (Fig. 2BC). For in-frame deletions, we modified a previously described protocol (Jackson et al., 2017). Briefly, this consists of creating the sequence of the deletion variant of interest, creating a homology model using the original crystal structure (PDB: 3V42) as the template, relaxing this model with Rosetta, and calculating the energy difference (ΔΔG) to the original structure (see Methods). This approach models both the effect of losing the side chain and backbone, but also the strain on the overall structure from the deletion (Fig. 2D). All data are included in the supplemental material (supplemental file 1). Data for the selected variants are included in Table 1.

Similar to previous studies (Abildgaard et al., 2019; Mathiassen et al., 2015; Matreyek et al., 2018; Nielsen et al., 2017; Scheller et al., 2019) we observe good correlation between predicted ΔΔGs and experimentally determined protein levels. Notably, both missense variants and in-frame deletions can be well-tolerated but can also lead to substantial loss of stability and degradation (Fig. 2B). In addition to the disease-associated variants in our dataset, we tested several additional deletion and missense variants, including two predicted and confirmed stable in-frame deletions ΔE455 and ΔA474 (Fig. 2E). In contrast, K508A shows substantially reduced stability, confirming the importance of the Lys side chain at position 508 (Fig. 2D). The ΔE509 variant was predicted to be as detrimental as its neighbours ΔK508 and ΔE510, however, it is only moderately destabilised (Fig. 2E). We speculate that this may be because the side chain points into solvent, where it is less critical than in the core of the folded protein, although this is not perfectly captured by our computational model. Lastly, we compared the predicted ΔΔGs with the variant allele frequency as reported in gnomAD (Karczewski et al., 2019). Similar to previously analysed proteins (Abildgaard et al., 2019; Nielsen et al., 2017; Scheller et al., 2019) this shows a “fishtail” distribution where the more common variants such as V400I and K508R have near-neutral stability changes, while some of the rarest and pathogenic variants are destabilised (Fig. 2F).

### Several FLCN variants are insoluble and mislocalized

Supported by the thermodynamic predictions, the reduced steady-state levels and increased proteasomal turnover suggest that the FLCN variants are misfolded and become PQC targets. In some cases, protein misfolding results in the formation of insoluble aggregates (Le Goff et al., 2016; Mathiassen et al., 2015; Shiber et al., 2013). To test this for the FLCN variants, we separated crude cell lysates into soluble (supernatant) and insoluble (pellet) fractions by centrifugation and analyzed the abundance of the FLCN variants in these fractions by Western blotting. Wild-type FLCN was found both in the soluble fraction and, presumably by interaction with membranes, in the pellet (Fig. S2AB). Several of the FLCN variants, particularly the R239C, ΔK508 and ΔE510 variants, were mostly insoluble (Fig. S2AB), suggesting that they are misfolded and may form aggregates.

Next, we proceeded to directly assess the subcellular localization of the FLCN variants by fluorescence microscopy. Consistent with other reports (Baba et al., 2008; Hasumi et al., 2008; Takagi et al., 2008), wild-type FLCN was evenly distributed throughout the cytosol and nucleus (Fig. 3A). This was also the case for the R239C, R362C, V400I and K508R variants. However, the nuclear localization of the other FLCN variants was strongly reduced, and in case of the ΔE510 variant, most of the cells (~70 %) displayed several perinuclear aggregate-like structures (Fig. 3A). Accordingly, retention of the ΔE510 variant was also apparent in filter trap assays (Fig. 3BC), where aggregated proteins fail to migrate through a nitrocellulose membrane (Westhoff et al., 2005).

**Fig. 3.**
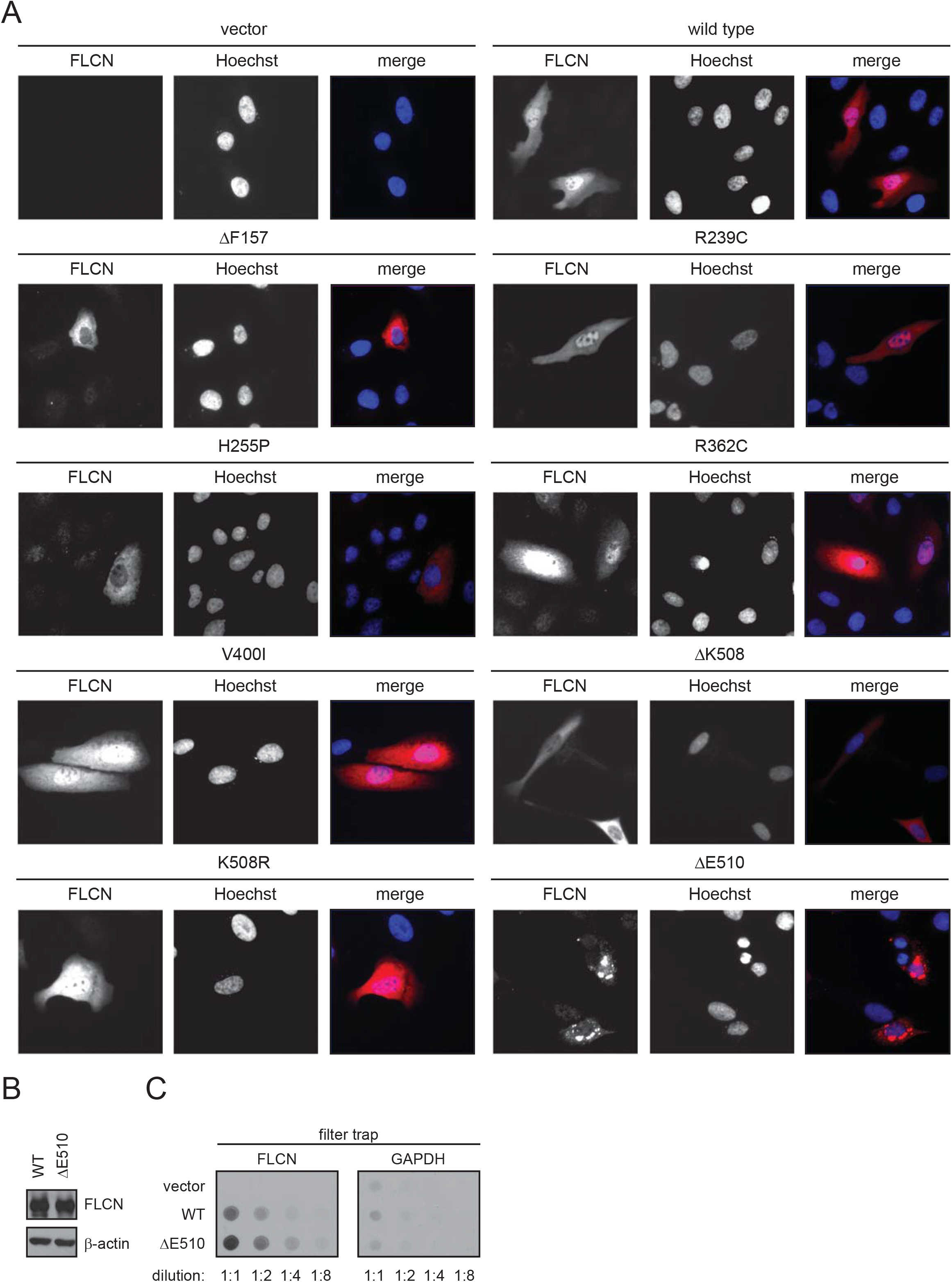
Certain BHD-linked FLCN variants are mislocalized. (A) The localization of the FLCN variants in transiently transfected U2OS cells was determined using antibodies to the 6His-tag. Hoechst was used to stain the nucleus. (B) As shown by SDS-PAGE and Western blotting of whole cell lysates, equal steady-state levels of wild-type (WT) FLCN and the ΔE510 variant were achieved by diluting the amount of WT FLCN plasmid with empty vector prior to transfection. (C) Dilution series of the whole cell lysates shown in (B) were filtered on a nitrocellulose membrane with a pore size of 0.2 μm and developed as a Western blot using antibodies to FLCN and, as a control, to GAPDH.

The absence of nuclear-localized FLCN has been observed for an unstable FLCN splice variant before (Bartram et al., 2017), and considering the unstable nature of these proteins, this could suggest that the observed increased protein turnover is restricted to the pool of nuclear FLCN. However, repeating the localization experiments in the presence of bortezomib did not result in any increased nuclear localization of the unstable variants or did otherwise affect the subcellular distribution of the FLCN variants (Fig. S3). This indicates that the lack of nuclear signal is caused primarily by a failure in nuclear import of FLCN rather than increased specific degradation of nuclear FLCN.

### FLCN binding protects FNIP1 and FNIP2 from proteasomal degradation

Genetic and structural studies have shown that FLCN function is intimately linked to its binding partners FNIP1 and FNIP2 (Schmidt and Linehan, 2018; Shen et al., 2019), and we therefore investigated the interplay between the FLCN variants and FNIP1/2. Upon introducing HA-tagged FNIP1 or FNIP2 expression constructs, we observed very low FNIP1/2 levels (Fig. 4AB). Since the FNIP1/2 levels increased in response to the proteasome inhibitor bortezomib, the low steady-state levels of FNIP1/2 are caused by proteasomal degradation (Fig. 4AB). However, upon co-expression of wild-type FLCN, the FNIP1/2 proteins were dramatically stabilized (Fig. 4AB), suggesting that FLCN protects FNIP1/2 from degradation. Indeed, when following the amount of FNIP1 and FNIP2 in cycloheximide-treated cultures (Fig. 4CD), the FNIP1/2 proteins were rapidly degraded (t_½_ ~ 16 hours), but stabilized (t_½_ ≫ 16 hours) by co-expression of FLCN (Fig. 4CDEF). Co-expression of the stable BHD-linked FLCN variants V400I and K508R, and to a lesser extent R239C and R362C, also led to FNIP1/2 stabilization, while this was less apparent for the unstable FLCN variants (Fig. 4GH).

**Fig. 4.**
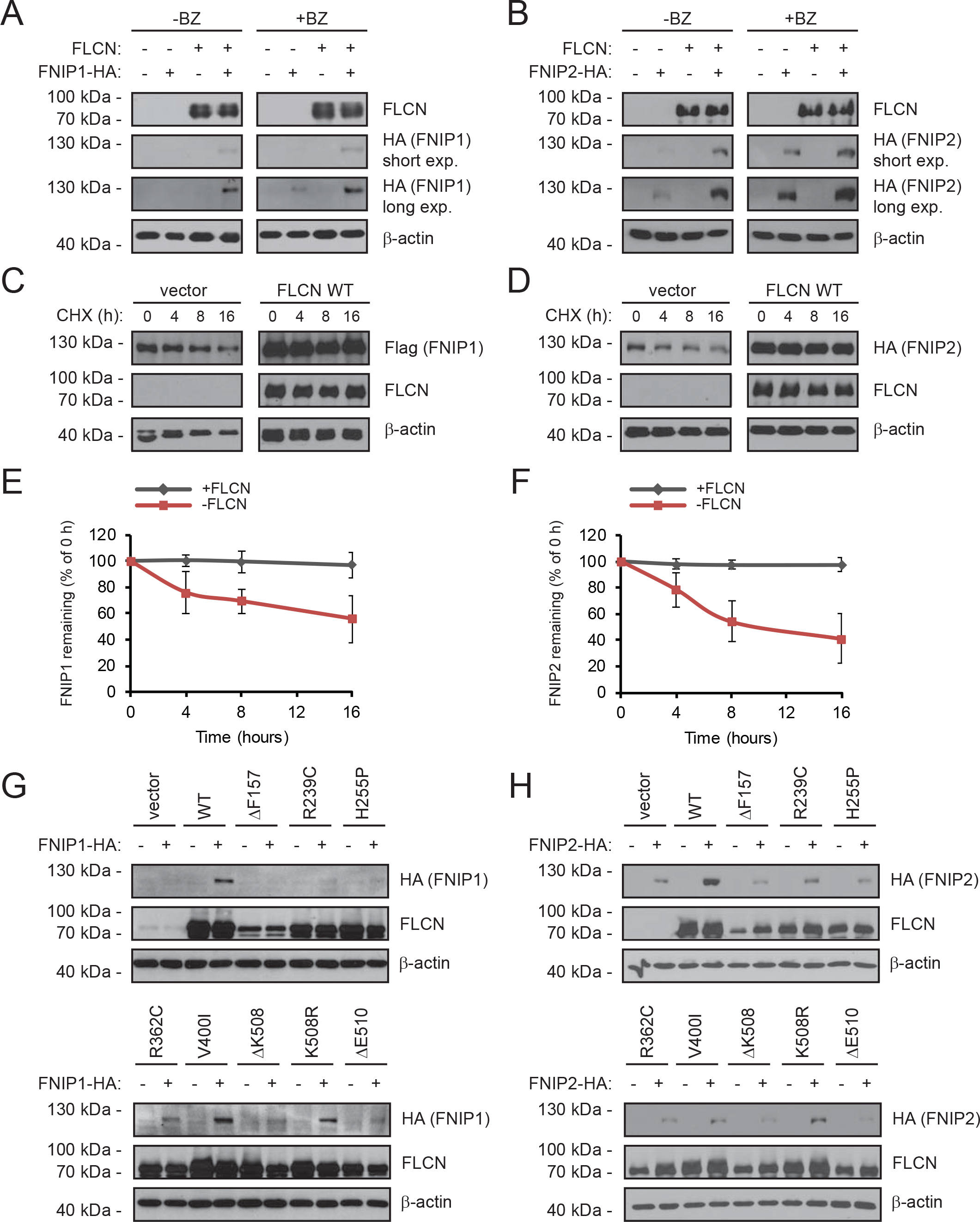
FLCN stabilizes FNIP1 and FNIP2. (A) U2OS cells transiently transfected to express either FLCN, HA-tagged FNIP1, or both FLCN and HA-FNIP1, were either untreated or treated with the proteasome inhibitor bortezomib (BZ) for 8 hours. Then the levels of FLCN and FNIP1 were compared by SDS-PAGE and Western blotting with antibodies to FLCN or the HA-tag on FNIP1. β-actin served as a loading control. (B) U2OS cells transiently transfected to express either FLCN, HA-tagged FNIP2, or both FLCN and HA-FNIP2, were either untreated or treated with the bortezomib. Then the levels of FLCN and FNIP2 were compared by SDS-PAGE and Western blotting with antibodies to FLCN or the HA-tag on FNIP2. β-actin served as a loading control. (C) The degradation of FNIP1 in the presence or absence of cotransfected FLCN was followed in cultures treated with the translation inhibitor cycloheximide (CHX) by SDS-PAGE and Western blotting with antibodies to the Flag-tag on FNIP1. FLCN and β-actin served as controls. (D) The degradation of FNIP2 in the presence or absence of co-transfected FLCN was followed in cultures treated with the translation inhibitor cycloheximide (CHX). (E) Quantification of blots as shown in (C) by densitometry normalized to the level at 0 hours. The error bars show the standard deviation (n = 3). (F) Quantification of blots as shown in (D) by densitometry normalized to the level at 0 hours. The error bars show the standard deviation (n = 3). (G) The levels of FLCN and FNIP1 were compared by SDS-PAGE and Western blotting in U2OS cells transiently transfected to express HA-tagged FNIP1 and the indicated FLCN variants. FLCN was detected with antibodies to FLCN. FNIP1 was detected using antibodies to the HA-tag. (H) The levels of FLCN and FNIP2 were compared by SDS-PAGE and Western blotting in U2OS cells transiently transfected to express HA-tagged FNIP2 and the indicated FLCN variants. FLCN was detected using antibodies to FLCN. FNIP2 was detected using antibodies to the HA-tag. In all panels β-actin served as a loading control.

Based on these data, we reasoned that FNIP1 and FNIP2 are protected from proteasomal degradation upon FLCN binding. Accordingly, in co-precipitation experiments with FNIP1/2 the V400I, K508R, R239C, and R362C FLCN variants all appeared adept at FLCN interaction, while the FNIP1/2 binding was strongly reduced for the other FLCN variants (Fig. 5AB). Co-expression of FNIP1 and FNIP2 did not lead to any mutual stabilization (Fig. 5C), excluding that that FNIP stabilization by FLCN occurs indirectly e.g. via a FNIP1-FNIP2 interaction.

**Fig. 5.**
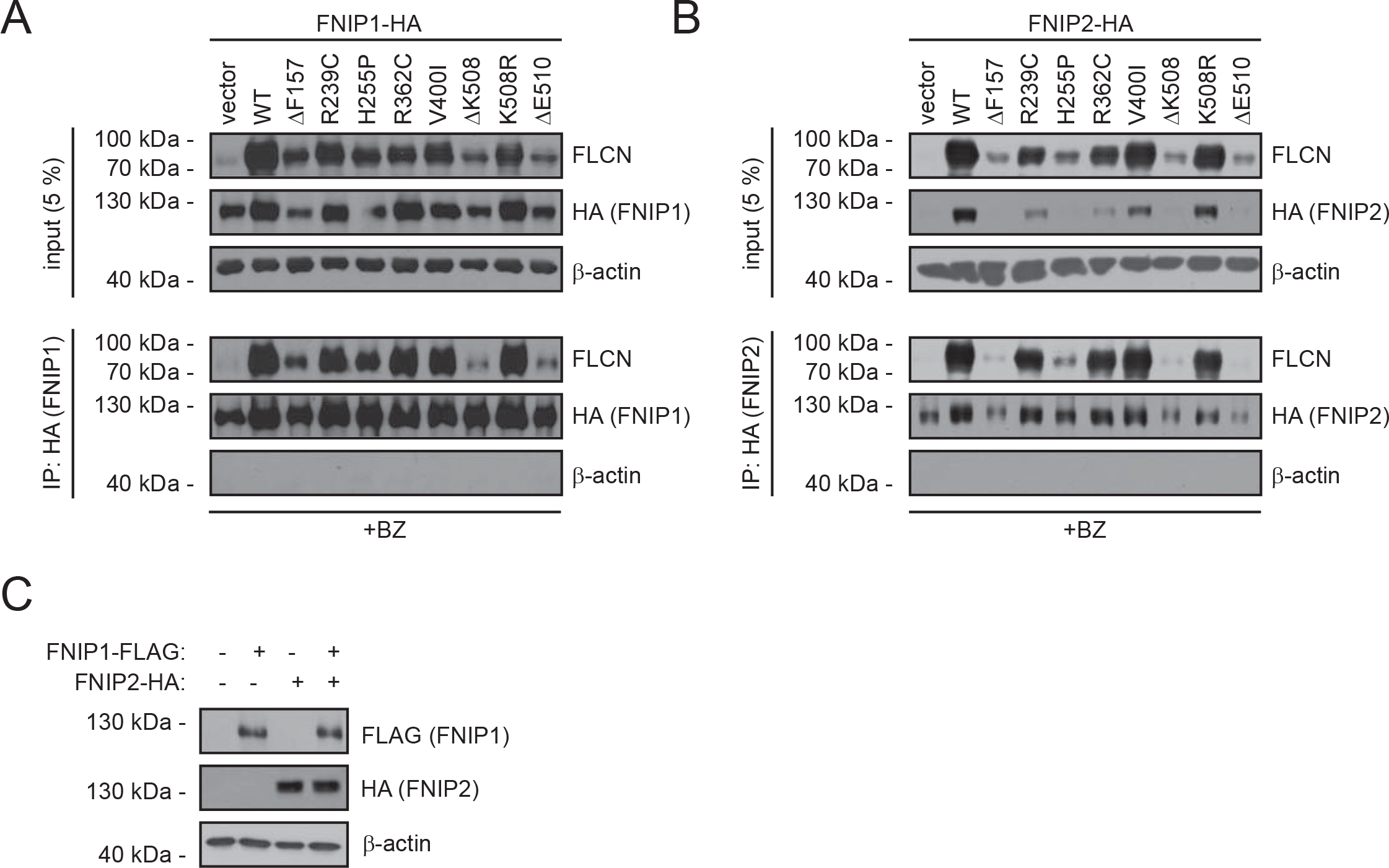
Some FLCN variants display reduced FNIP1/2 binding. (A) The FLCN variants were analyzed for interaction with FNIP1. HA-tagged FNIP1 was immunoprecipitated and analyzed by blotting. Input samples (5%) were included as a control. To obtain an adequate amount of the FNIP1 and FLCN proteins, the cells were treated with bortezomib, for 8 hours before harvest. (B) The FLCN variants were analyzed for interaction with FNIP2. HA-tagged FNIP2 was immunoprecipitated and analyzed by blotting. Input samples (5%) were included as a control. Similar to above, the cells were treated for 8 hours with bortezomib (BZ) prior to harvesting and lysis. (C) U2OS cells, transiently transfected to express either Flag-tagged FNIP1, HA-tagged FNIP2, or both, were analyzed by blotting.

### A yeast screen links FLCN turnover to chaperones and the deubiquitylating enzyme Ubp15/USP7

To further analyze the degradation of the unstable FLCN variants we took advantage of a yeast selection system for protein stability (Geffen et al., 2016). Specifically, we expressed wild-type FLCN (as a control) and the unstable ΔE510 variant as fusion proteins with the orotidine-5’-phosphate (OMP) decarboxylase Ura3 enzyme in a strain carrying a deletion of the endogenous *URA3* gene (*ura3*Δ) (Fig. 6A). Since proteasomal protein degradation is processive (Bard et al., 2018), degradation of the ΔE510 fusion protein, should lead to reduced amounts of the Ura3 fusion partner. Accordingly, we observed the FLCN ΔE510 fusion protein was rapidly degraded via the proteasome (Fig. 6B), and the ΔE510 strain displayed a reduced growth in media lacking uracil (Fig. 6C). As expected the growth defect was completely suppressed by addition of sublethal amounts of the proteasome inhibitor bortezomib (Fig. 6C), and we therefore continued to screen for components targeting the unstable FLCN variants for degradation. To this end, the ΔE510 yeast expression construct was introduced into a library of 4992 individual gene deletion mutants (Winzeler et al., 1999), one-by-one, by mating. After sporulation and selection of haploid cells with the gene deletion (G418 resistant) and the Ura3 reporter fusion construct, the resulting haploids were scored for growth by monitoring the colony sizes on solid media lacking uracil. In total 27 mutants caused significantly increased growth, suggesting a stabilization of the ΔE510 variant, while 135 mutants displayed a significantly reduced growth, indicating that the ΔE510 variant was less abundant in these cells. All 162 significant hits from this screen are included in the supplemental material (supplemental file 2). To test the validity of the screen we selected two hits, *ydj1*Δ and *ubp15*Δ, for further analyses. In agreement with the results from the high-throughput screen, introducing the ΔE510 fusion in *ydj1*Δ cells led to a visibly increased growth, while the *ubp15*Δ cells appeared to grow more poorly (Fig. 6D). This difference was also apparent in the steady-state levels of the ΔE510 fusion. Hence, in *ydj1*Δ cells we observed a higher level of ΔE510 fusion protein, while in the *ubp15*Δ strain, the ΔE510 protein level was reduced (Fig. 6E).

**Fig. 6.**
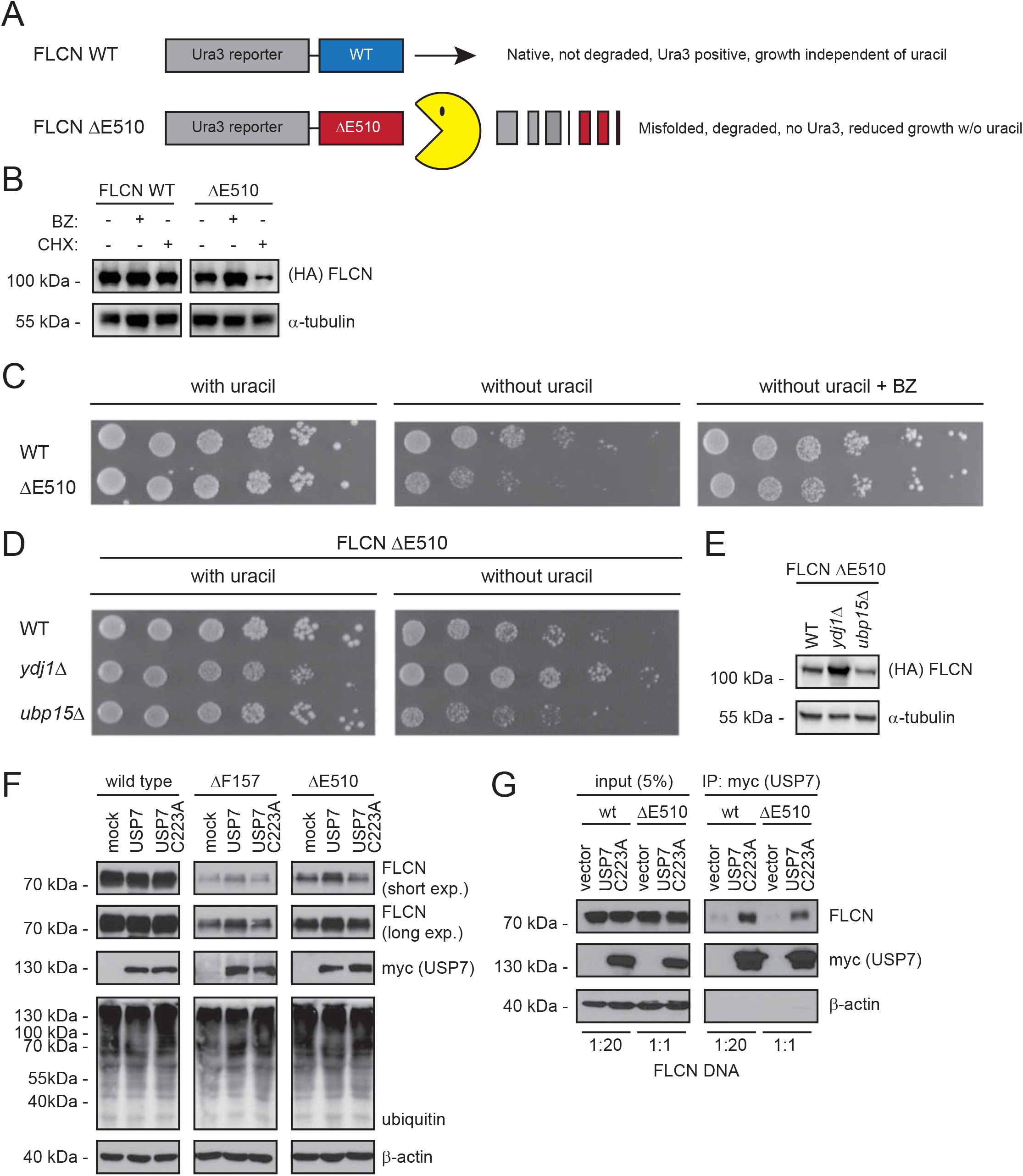
The USP7 DUB regulates FLCN protein levels. (A) Schematic illustration of the utilized yeast system for high-throughput scoring of FLCN protein stability. Briefly, either wild-type FLCN or the ΔE510 FLCN variant is fused to the C-terminus of a HA-tagged Ura3-reporter protein. In case of the ΔE510 FLCN variant, we expect the protein to be destabilized and degraded (Pac-Man), leading to reduced levels of the Ura3 protein and thereby a decreased growth in the absence of uracil. (B) The amount of FLCN and FLCN ΔE510 Ura3-fusion protein was analyzed in cultures treated with (+) or without (-) 1 mM of the proteasome inhibitor bortezomib (BZ) for 3 hours, or where protein synthesis was inhibited with 100 μg/ml of cycloheximide (CHX) for 4 hours. For detection on Western blots antibodies to HA (to detect FLCN), and, as a loading control, to α-tubulin, were used. (C) Growth of wild-type yeast cells transformed with Ura3-fusion vectors for wild-type (wt) FLCN or FLCN ΔE510 was compared by spotting serial dilutions on solid media with uracil (left panel), without uracil (central panel), or without uracil, but supplemented with bortezomib (BZ) (right panel). (D) Growth wild-type (WT), *ydj1*Δ and *ubp15*Δ yeast cells transformed with the Ura3-fusion vector for FLCN ΔE510 was compared by spotting serial dilutions on solid media with uracil (left panel) and without uracil (right panel). (E) The level of the FLCN ΔE510 Ura3-fusion protein in lysates was compared by blotting using antibodies to HA (to detect FLCN), and, as a loading control, to α-tubulin. (F) FLCN variants were transfected into U2OS cells with an expression vector for either wild-type myc-tagged USP7 or the catalytically dead USP7 C223A variant. Lysates were analyzed by SDS-PAGE and Western blotting using antibodies to FLCN, myc (to detect the co-expressed USP7) and ubiquitin. β-actin served as a loading control. (G) The wild-type and ΔE510 FLCN variants were analyzed for interaction with the catalytically dead USP7 C223A variant. Myc-tagged USP7 was immunoprecipitated using the myc-trap system and analyzed by blotting. Input samples (5%) were included as a control. To achieve equal amounts of the FLCN proteins, the cells were transfected with 20-fold less wild-type FLCN DNA compared to FLCN ΔE510 DNA.

Ydj1 is a DnaJ domain family co-chaperone linked to proteasomal degradation of misfolded chaperone clients (Maurer et al., 2016), while Ubp15 is a deubiquitylating enzyme which counteracts E3s to block degradation. Whereas human cells encode a number of J-domain proteins that are homologous to the yeast Ydj1, the yeast Ubp15 is in human cells represented by a single orthologue called USP7. Next, we therefore co-transfected U2OS cells to express FLCN and either wild-type USP7 or, as a control, a catalytically dead USP7 variant carrying a C223A substitution of the active site cysteine residue. We note that the overexpression of USP7 did not affect the overall ubiquitylation of cell proteins (Fig. 6F). In agreement with the results in yeast cells, overexpression of the wild-type USP7, but not of the catalytically dead USP7 variant, led to a stabilization of the FLCN ΔF157 and ΔE510 variants (Fig. 6F), suggesting that USP7 deubiquitylates these proteins.

Since we found that FLCN contains multiple potential TRAF- and UBL1/2-domain binding consensus motifs (Fig. S4 & S5) that could mediate direct interaction to USP7 (Kim and Sixma, 2017), we tested if FLCN and USP7 interact by co-precipitation. To achieve equal amounts of wild type and ΔE510, we transfected the U2OS cells with 20-fold less plasmid encoding wild-type FLCN compared to ΔE510, but equal amounts of a plasmid encoding myc-tagged USP7 C223A. We observed a clear interaction between USP7 and FLCN (Fig. 6G), supporting further a role of USP7 in regulating FLCN turnover.

### Effect of temperature on FLCN stability and function

In some cases, PQC targets can be stabilized structurally by lowering the temperature, which in turn leads to a reduced degradation and increased steady-state level (Nielsen et al., 2017). To test this for the FLCN variants, we compared their steady-state levels at 37 °C and 29 °C. Importantly, lowering the temperature to 29 °C does not affect the ubiquitin-proteasome system in U2OS cells (Fig. S6) (Nielsen et al., 2017). Several of the FLCN variants were stabilized at 29 °C (Fig. 7A), indicating that these variants are indeed structurally stabilized at the lower temperature. This effect was most apparent for the R362C variant (Fig. 7A), in line with our observation that this variant is borderline stable already at 37 °C, and interacts with and partially stabilizes FNIP1/2 (Table 1). Thus, a slight structural stabilization conveyed by lowering the temperature is enough for this variant to evade the PQC-mediated clearance. Accordingly, we reasoned that perhaps the tumor suppressor function of some FLCN variants could be restored upon lowering the temperature. We therefore investigated the ability of the FLCN variants to suppress the growth of *FLCN*-negative FTC-133 cells suspended in soft agar at 37 °C and 29 °C. The FLCN expression constructs were transfected into FTC-133 cells as stable mixed cell populations, and after four weeks at 37 °C or 29 °C colonies were counted. Wildtype FLCN led to roughly 50% reduction in colony formation at both temperatures (Fig. 7B). The stable variants, V400I and K508R, both behaved like wild type and suppressed growth to a level similar to wild-type FLCN (Fig. 7B). In line with a stabilization at low temperature, the R362C variant significantly reduced colony formation only at 29 °C (Fig. 7B), which was not apparent for any of the other FLCN variants.

**Fig. 7.**
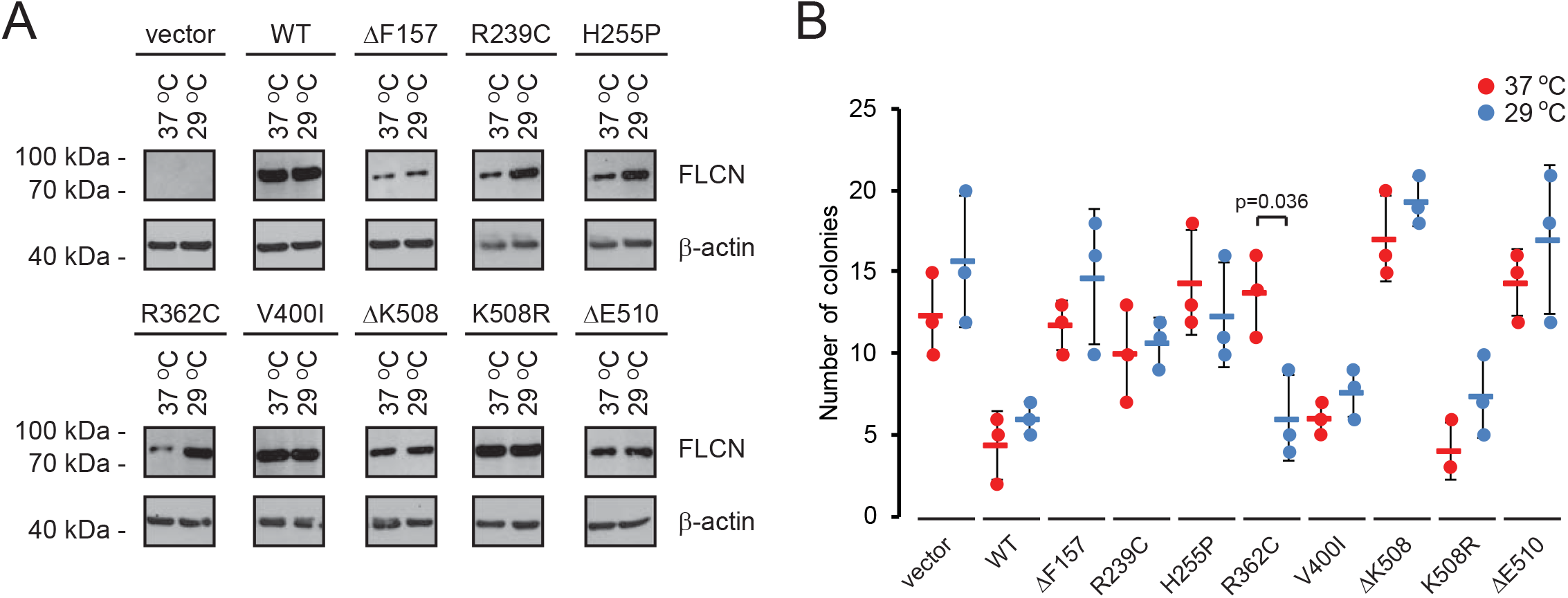
BHD-linked FLCN variants are stabilized at reduced temperatures. (A) The steady-state level of wild-type (WT) FLCN and the selected FLCN variants, expressed in U2OS cells, was compared at 37 °C and 29 °C. FLCN was detected using antibodies to FLCN. β-actin served as a loading control. (B) Colony formation at 37 °C (red) and 29 °C (blue) of FTC-133 cells transfected with the indicated expression constructs. The bars indicate the standard deviation (n = 3, p-value based on students t-test).

## Discussion

Certainly, point mutations may directly ablate protein function without grossly affecting protein structure, e.g. by disrupting the active site of an enzyme. More commonly, however, missense mutations or in-frame deletions result in a structural perturbation or destabilization of the encoded protein (Stein et al., 2019), and indeed a previous report has shown that several BHD-linked single site FLCN variants are destabilized (Nahorski et al., 2011). The results presented here are perfectly in line with this observation, and show that the structural destabilization, conferred by the mutations, leads to rapid proteasomal degradation of the FLCN variants, and in the rare cases that BHD syndrome is triggered by such single site mutations, the disease should be considered a protein misfolding disease.

We found that some of the FLCN variants are mislocalized in cells and one variant (ΔE510) formed perinuclear protein aggregates, suggesting that at least this variant is highly misfolded. Often, aggregated proteins are cleared by autophagy. However, similar to aggregation-prone variants of VHL (Le Goff et al., 2016), ataxin-3 (Chai et al., 1999) and DJ-1 (Mathiassen et al., 2015), the FLCN ΔE510 protein appeared to be entirely degraded via the proteasome, which suggests that molecular chaperones may dissolve the ΔE510 aggregates prior to degradation.

Our screen in yeast cells revealed two oppositely acting regulators of FLCN stability. In agreement with chaperones playing a role in FLCN turnover, deletion of the Ydj1 co-chaperone led to a stabilization of the FLCN ΔE150 variant. Deletion of the Ubp15, on the other hand, destabilized the FLCN ΔE510 variant. This observation fits with Ubp15/USP7 possessing deubiquitylating activity, which by catalyzing the removal of ubiquitin chains from proteins destined for the proteasome counters the degradation. Previous studies have identified numerous binding partners and substrates of USP7 (Kim and Sixma, 2017; Sowa et al., 2009), including p53, p16^INK4a^ and WASH (Cummins et al., 2004; Hao et al., 2015; Li et al., 2004; Maertens et al., 2010). Typically, these interact with USP7 through its TRAF domain and/or the UBL1/2 domains, and consensus motifs in USP7-binding partners have been described (Kim and Sixma, 2017). We note that FLCN contains multiple of these potential TRAF- and UBL1/2-domain binding consensus motifs, which could mediate direct interaction to USP7. However, notably USP7 appears to preferentially interact with E3 ubiquitin-protein ligases (Kim and Sixma, 2017), as exemplified by the effect of USP7 on p53. Hence, in case of p53, USP7 has been shown to deubiquitylate and stabilize the E3 MDM2, which in turn leads to an increased degradation of p53 (Cummins et al., 2004; Li et al., 2004; Reverdy et al., 2012; Turnbull et al., 2017). Presumably since PQC E3s display overlapping substrate specificity and are highly redundant (Samant et al., 2018; Theodoraki et al., 2012) our screen failed to uncover the E3(s) that target FLCN for degradation, and our results on USP7 suggest that USP7 directly deubiquitylates FLCN, rather than targeting an FLCN-specific E3.

Classically molecular chaperones are viewed as protein folding machines that capture non-native proteins and, through cycles of binding and release, catalyze folding into the correct native state. However, chaperones also directly interact and collaborate with components of the ubiquitin-proteasome system (Gowda et al., 2013; Kandasamy and Andreasson, 2018; Kriegenburg et al., 2012; Kriegenburg et al., 2014; Maurer et al., 2016; Samant et al., 2018). For instance, HSP70 and HSP90 directly associate with the E3 ubiquitin ligase, CHIP, which in turn ubiquitylates the chaperone clients to ensure their degradation (Connell et al., 2001; Esser et al., 2004). Recently, it was shown that also wild-type FLCN is a HSP90 client protein and the FNIP1/2 binding partners function as cochaperones that facilitate FLCN folding and ensure FLCN stability (Sager et al., 2019; Woodford et al., 2016). In our assays, we observed that FLCN also functions to stabilize FNIP1/2, and this stabilization depends on FNIP1/2 interaction. Hence, for those FLCN variants that are misfolded and fail to interact with FNIP1/2, the FNIP1/2 levels are reduced, due to proteasomal degradation. Subunits in multiprotein assemblies are generally more stable once incorporated into a complex and unassembled (orphan) proteins, produced in excess, are targeted for degradation (McShane et al., 2016; Yanagitani et al., 2017). In the case of FLCN and the FNIP1/2 proteins, however, the balance in their stoichiometry might also be regulated by environmental cues. Indeed, it was recently shown that in the presence of nutrients, FNIP2 is phosphorylated by casein kinase 1 (CK1) and degraded by the proteasome (Nagashima et al., 2017). The experiments, presented here, were also performed under nutrient-rich conditions, and it is possible that the observed degradation of FNIP1/2 is triggered by CK1 phosphorylation of orphan FNIP1/2.

Intriguingly, we observed a clear correlation between the degradation rate of the unstable variants and several of the other tested parameters. Thus, all the unstable variants displayed reduced solubility, and with the exception of R239C and R362C, the unstable FLCN variants were all exclusively localized in the cytosol and failed to associate with and stabilize FNIP1 and FNIP2 (Table 1).

Of the tested FLCN variants, only two, V400I and K508R, behaved like wild-type FLCN with regard to level, turnover, interaction, localization and cell proliferation (Table 1). Since valine/isoleucine and lysine/arginine, are, respectively, chemically similar amino acids, it is not surprising that these substitutions do not render FLCN structurally unstable. However, importantly, these findings do not preclude that the V400I and K508R variants are pathogenic. According to the gnomAD database (Karczewski et al., 2019), the V400I variant is rare (~1:40,000), while K508R is more common (~1:5,000), suggesting that if the K508R mutation is pathogenic, it is likely to have a reduced penetrance (the estimated prevalence of BHD syndrome according to the Orphanet database is 1:200,000). ClinVar lists K508R as likely benign. However, a recent report has shown that although introducing the K508R variant in *Flcn* knock-out mice prolonged survival, the mice later developed cystic kidneys and succumbed to renal failure (Hasumi et al., 2017), indicating that K508R might be a weakly pathogenic mutation. It is possible that the slow onset, observed for the K508R mouse model, is connected with the K508R variant being only slightly more unstable than wild-type FLCN *in vivo*. However, it is more likely that any potential pathogenicity associated with the V400I and K508R variants, is not caused by structural destabilization, but instead associated with other properties of the FLCN protein that we did not test here.

Since we found that several of the variants were stabilized by lowering the temperature, these proteins are probably not highly destabilized or misfolded. Indeed, for the R362C variant, we observed an increased cellular function when the temperature was reduced. Thus, similar to other genetic diseases, including cystic fibrosis (Van et al., 2011) and von Hippel-Lindau disease (Yang et al., 2013), as well as certain cancer-linked mutations in p53 (Joerger and Fersht, 2016), it may be possible to stabilize some misfolded FLCN protein variants using a small molecule that either blocks the PQC system or directly stabilizes the FLCN protein structure (Kampmeyer et al., 2017b). However, to achieve this it is essential with further, in particular structural and biophysical, studies of both wild-type and disease-linked FLCN protein variants.

## Materials and methods

### Buffers

Buffer A: 50 mM Tris/HCl pH 7.4, 150 mM NaCl, 1 mM EDTA, 0.5 % NP-40, 1 mM PMSF and Complete protease inhibitors (Roche). Buffer B: 50 mM Tris/HCl pH 7.4, 1 mM EDTA, 150 mM NaCl, 1 mM PMSF and Complete protease inhibitors (Roche). PBS: 10 mM Na_2_HPO_4_, 1.8 mM KH_2_PO_4_, 137 mM NaCl, 3 mM KCl, pH 7.4.

### Plasmids

Full-length wild-type human *FLCN* cDNA carrying an N-terminal RGS6xHis-tag was expressed from pcDNA3.1 (Genscript). USP7 was expressed with an N-terminal myc-tag from pcDNA3.1 (Genscript). All mutations were generated by Genscript. The pcDNA3.1-FLAG-FNIP1 and the pcDNA3.1+N-eGFP-FLCNΔE510 vectors were generated by Genscript. The pIRES2-FLCN plasmids were kindly provided by Dr. E. R. Maher (Birmingham, UK). The pEGFP-N1-FLCN, pRK5-HA-FNIP1 and pRK5-HA-FNIP2 constructs were kindly provided by Dr. D. M. Sabatini (MIT, USA).

### Cell culture

U2OS cells (ATCC) were propagated in Dulbecco’s Modified Eagle medium (DMEM) with 10 % fetal-calf serum (Invitrogen), and supplemented with 5000 IU/mL penicillin, 5 mg/mL streptomycin and 2 mM glutamine at 37 °C. FTC-133 cells (ECACC) were similarly cultured but in DMEM:Ham’s F12 (1:1) medium with 2 mM glutamine and 10 % fetal-calf serum (Invitrogen). FugeneHD (Promega) was used for transfections following the manufacturer’s instructions. Cell imaging and quantifications were performed as described before (Nielsen et al., 2017). Protein aggregates were counted manually.

### Electrophoresis and blotting

SDS-PAGE was performed on 12.5 % acrylamide gels. For Western blotting 0.2 μm nitrocellulose membranes were used. For blocking, the membranes were incubated in 5 % fat-free milk powder and 0.1 % Tween-20 in PBS. Antibodies and their sources were: anti-ubiquitin (Z0458, DAKO), anti-Rpt6 (p45-110, Enzo Biosciences), anti-20S proteasome α-subunits (MCP231, Enzo Biosciences), anti-FLCN (D14G9, Cell Signaling Technology), anti-GFP (3H9, Chromotek), anti-myc (9E1, Chromotek) anti-HA (12CA5, Roche), anti-Flag (M2, Sigma), anti-RGSHis (34650, Qiagen), anti-β-actin (AC74, Sigma), anti-NaK-ATPase (C464.6, Sigma), anti-LC3A/B (D3U4C, Cell Signaling Technology), anti-GAPDH (14C10, Cell Signaling Technology), anti-α tubulin (TAT-1, 00020911 Sigma). All secondary antibodies were purchased from DAKO. The Un-Scan-It v6.1 software (Silk Scientific) was used for densitometry.

### Protein degradation experiments

Bortezomib (LC Laboratories) and chloroquine (Sigma) were used at 15 μM and 20 μM respectively, for 8 hours in serum-free media. Degradation was followed in cultures treated with cycloheximide (CHX) as described previously (Nielsen et al., 2017).

### Co-precipitation experiments

Binding studies were performed as described before (Nielsen et al., 2017), using anti-HA resin (Sigma) or myc-trap beads (Chromotek). Bound proteins were eluted directly in SDS sample buffer.

### FLCN solubility

To estimate the solubility of the FLCN variants, a confluent 6-well dish of transfected U2OS cells was lysed in 200 μL buffer B by three 20 second rounds of sonication on ice. The lysate was then centrifuged at 15000 g for 30 minutes at 4 °C. Finally, 4x concentrated SDS sample buffer was added to the supernatant to obtain a 1x SDS sample buffer lysate. The pellet was resuspended in 1x SDS sample buffer to match the volume of the supernatant.

### Filter trap assays

Filter trap assays were performed largely as described previously (Westhoff et al., 2005). Briefly, to achieve equal cellular expression of wild-type FLCN and FLCN Δ510 protein, the plasmid encoding wild-type FLCN was diluted with empty control vector prior to transfection. Transfected cells were harvested in ice-cold PBS using a cell scraper and lysed by sonication for three times 10 seconds in ice-cold PBS containing Complete protease inhibitors (Roche). Unbroken cells were removed by centrifugation at 500 g for 2 minutes. SDS was added to a final concentration of 2 % and three twofold serial dilutions in 2 % SDS in PBS were prepared. Samples of 200 μL were loaded onto a 0.2 μm nitrocellulose membrane affixed in a BioRad Bio-Dot apparatus (BioRad) pre-equilibrated with 100 μL PBS per well. The wells were then washed once with 400 μL 2 % SDS in PBS and then once with 200 μL 2 % SDS in PBS. The nitrocellulose membrane was then removed from the apparatus and washed briefly in PBS, before blocking and development as a Western blot using antibodies to FLCN and as a control GAPDH.

### Soft agar colony formation assays

Soft agar colony formation assays were performed as described (Nahorski et al., 2011) using pools of FTC-133 cells stably transfected to express the FLCN variants. After four weeks, colonies greater than 100 μm in diameter were counted. The experiments were performed in triplicates.

### Yeast strains and techniques

All yeast strains were from the Euroscarf collection and transformations were performed using lithium acetate (Gietz and Woods, 2002). For analyzing growth on solid media, exponential phase cultures were diluted to an OD_600nm_ of 0.40 and 5 μL of 5-fold serial dilutions were applied in spots on agar plates. Cell extracts for Western blotting were prepared with glass beads in trichloroacetic acid (Sigma) as described (Cox et al., 1997). Cycloheximide (CHX) (Sigma) was used at a concentration of 100 μg/mL. Bortezomib (BZ) (LC Laboratories) was used at a concentration of 1 mM. The screening of the yeast gene deletion library (Winzeler et al., 1999) was performed by first transforming the FLCN-Ura3 reporter plasmid (pTR1412-FLCNΔE510) into the query strain Y7092 (*MATα can1*Δ*::STE2pr-Sp_his5 lyp1*Δ *ura3*Δ*0 leu2*Δ*0 his3*Δ*1 met15*Δ*0 geneX::KanMX6*) and selecting transformants on SC-Leu medium. Next, the transformed strain was mated to the yeast gene deletion strain collection by automated pinning (RoToR, Singer Instruments, UK). Selection for diploids, sporulation, and selection for haploid progeny containing the gene deletion (G418^R^) and the FLCN-Ura3 reporter plasmid was performed essentially as described (Tong and Boone, 2007).

### Calculation of ΔΔGs for missense variants

We used the Rosetta cartesian_ddg protocol (Park et al., 2016) and the high-resolution structure of the C-terminal FLCN domain (PDB 3V42) (Nookala et al., 2012) to calculate ΔΔGs for missense variants, as also previously described (Scheller et al., 2019). Three iterations were performed for each of the 20 possible amino acid variants at each position, including the wild type for reference. The value reported here is the difference between the average of the wild-type Rosetta energies and the average for the respective variant. All scores are divided by 2.9 to bring them to a kcal/mol scale (Frank DiMaio, personal communication).

### ΔΔGs for in-frame deletion variants

First 25 homology models were created (Webb and Sali, 2016) for each single-amino-acid-deletion in the C-terminal domain, using PDB 3V42 (Nookala et al., 2012) as the template. Each homology model was relaxed in two independent trajectories using the ”/path/to/rosetta/source/bin/relax.linuxgccrelease -database /path/to/rosetta/database -l list_w_models.lst -ignore_unrecognized_res -use_input_sc -constrain_relax_to_start_coords - flip_HNQ -no_optH false -relax:fast -nstruct 2” command line, based on Jackson et al. (Jackson et al., 2017). The median of the resulting 50 scores was calculated to estimate the ΔG (in Rosetta Energy Units) of the respective deletion variant. To estimate ΔΔGs, the 10^th^ percentile of the ΔG distribution was subtracted. As for the missense variant ΔΔGs above, scores were divided by 2.9 to bring them to a scale that corresponds to kcal/mol.

## Acknowledgements

The authors thank Dr. Thomas Hansen, Dr. Elin Pietras, Dr. Cornelia Steinhauer, Rasmus Scheller, Sven Larsen-Ledet and Anne-Marie Lauridsen for excellent technical assistance, and Dr. Thomas van Overeem Hansen and Dr. Klavs B. Hendil for helpful discussions and comments on the manuscript. We acknowledge Dr. Elena Papaleo, who contributed to early parts of this project. We thank Dr. Charlie Boone for sharing reagents. Expression plasmids were kindly provided by Dr. David M. Sabatini and Dr. Eamonn R. Maher.

## Author contributions

L.C., A.S., M.G.T., S.V.N., L.N., and C.S. conducted the experiments. L.C., A.S., S.V.N., M.L., T.R., K.L.L., and R.H.P. designed the experiments and analyzed the data. L.C., A.S., K.L.L., and R.H.P. wrote the paper.

## Supplemental material

Supplemental figures and references, SupplementalFigures.pdf

Supplemental file 1, In silico saturation mutagenesis data, SupplementalFile1.xlsx

Supplemental file 2, Results from the yeast screen, SupplementalFile2.xlsx

