## Supplemental figures for "Folliculin variants linked to Birt-Hogg-Dubé syndrome are targeted for proteasomal degradation"

### *Supplemental material*


| | |
|---|---|
| *Supplemental figure Fig.S1 (The FLCN variants are not autophagy targets.):* | *p2* |
| *Supplemental figure Fig. S2 (Several BHD-linked FLCN variants are insoluble)* | *p3* |
| *Supplemental figure Fig.S3 (The subcellular localization of FLCN):* | *p4* |
| *Supplemental figure Fig.S4 (FLCN contains multiple potential USP7 interaction motifs):* | *p5* |
| *Supplemental figure Fig.S5 (The potential USP7 binding sites in the FLCN structure):* | *p6* |
| *Supplemental figure Fig.S6 (The ubiquitin-proteasome system at 29 °C):* | *p7* |
| *Supplemental references:* | *p8* |

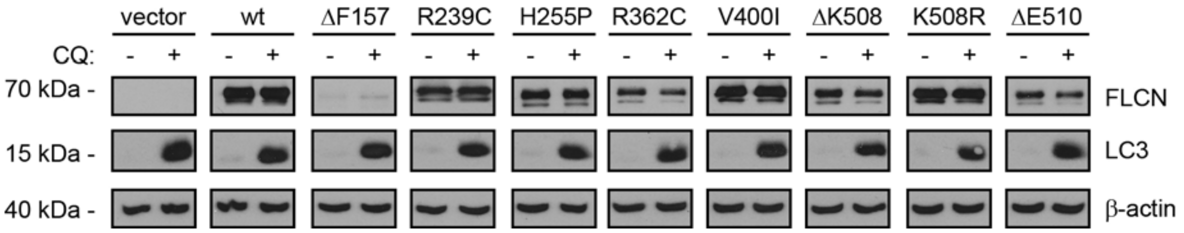


**Figure S1** *The FLCN variants are not autophagy targets.*
The steady-state levels of the FLCN variants were compared by Western blotting, using antibodies to FLCN, in cultures that were either untreated or treated with the autophagy inhibitor chloroquine (CQ) for 8 hours. β-actin served as a loading control, while blotting for the autophagy substrate, LC3, was included as a control for successful inhibition of autophagy.

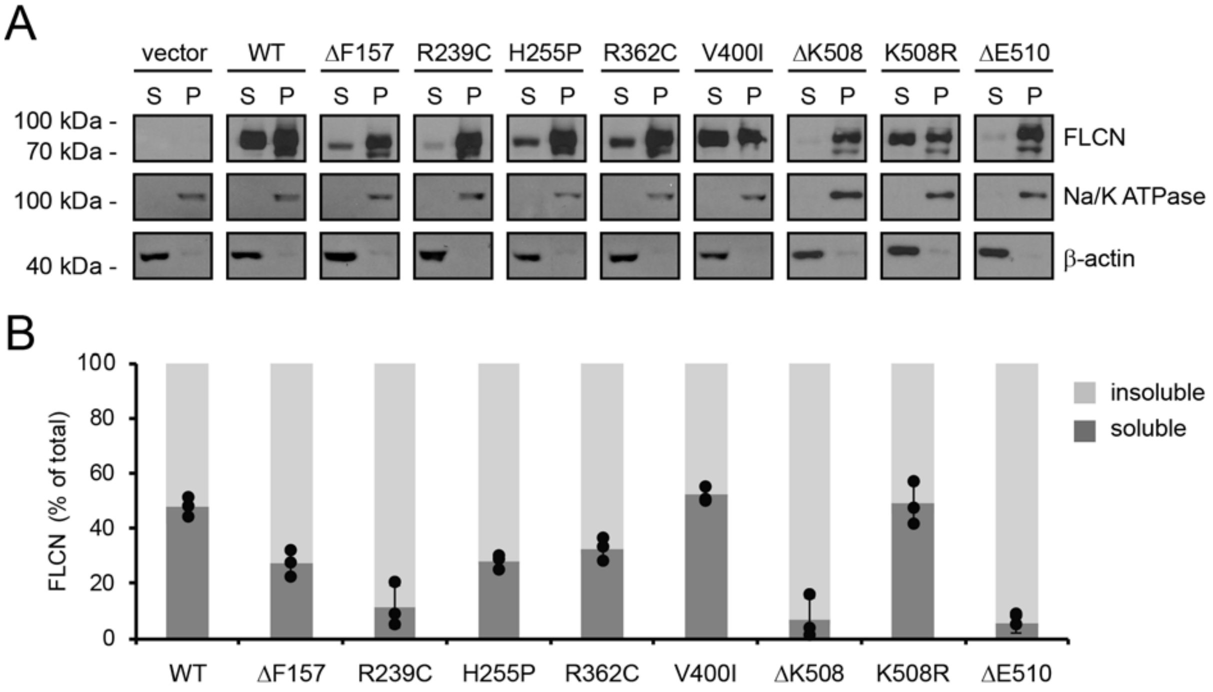


**Figure S2** *Several BHD-linked FLCN variants are insoluble.*
(A) The solubility of the selected FLCN variants. Samples of whole cell lysates were separated into a soluble supernatant (S) fraction and an insoluble pellet (P) fraction by centrifugation. FLCN concentrations were determined by SDS-PAGE and Western blotting with antibodies to FLCN. Na/K ATPase and β-actin were used as loading controls. (B) Quantification of blots as shown in (A) by densitometry. The soluble fractions are shown in dark grey, the insoluble pellet fractions in light grey. The error bars show the standard deviation (n = 3).

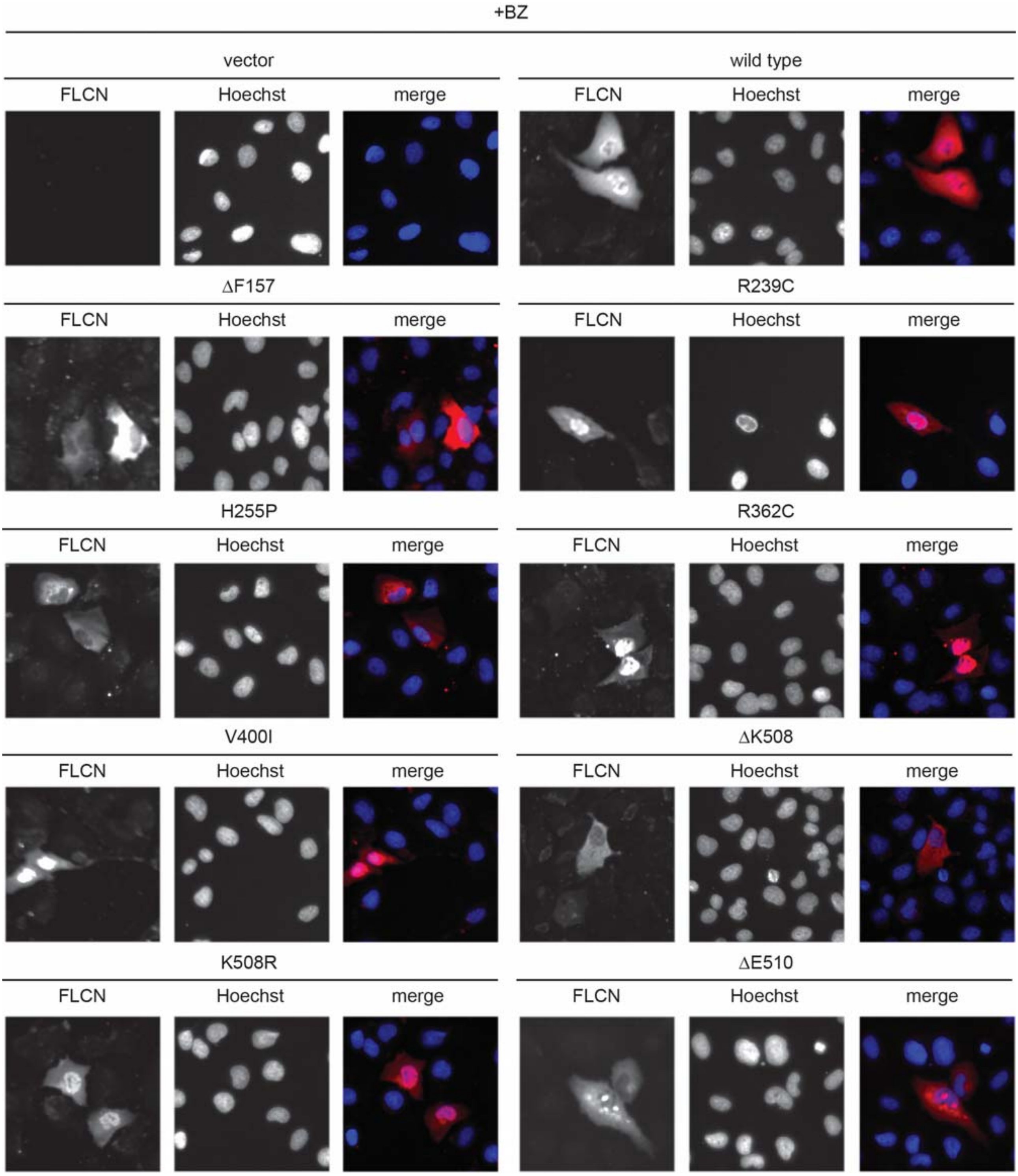


**Figure S3** *The subcellular localization of FLCN is unchanged by proteasome inhibition.*
U2OS cells transiently transfected to express 6His-tagged wild type FLCN and selected FLCN variants were treated with the proteasome inhibitor bortezomib (BZ) for 8 hours and analyzed by fluorescence microscopy. FLCN was stained using antibodies to the 6His-tag, and Hoechst was used to mark the nucleus.

```
MNAIVALCHFCELHGPRTLFCTEVLHAPLPQGDGNEDSPGQGEQAEEEEGGIQMNSRMRA  60
HSPAEGASVESSSPGPKKSDMCEGCRSLAAGHPGYISHDKETSIKYVSHQHPSHPQLFSI 120
VRQACVRSLSCEVCPGREGPIFFGDEQHGFVFSHTFFIKDSLARGFQRWYSIITIMMDRI 180
YLINSWPFLLGKVRGIIDELQGKALKVFEAEQFGCPQRAQRMNTAFTPFLHQRNGNAARS 240
LTSLTSDDNLWACLHTSFAWLLKACGSRLTEKLLEGAPTEDTLVQMEKLADLEEESESWD 300
NSEAEEEEKAPVLPESTEGRELTQGPAESSSLSGCGSWQPRKLPVFKSLRHMRQVLGAPS 360
FRMLAWHVLMGNQVIWKSRDVDLVQSAFEVLRTMLPVGCVRIIPYSSQYEEAYRCNFLGL 420
SPHVQIPPHVLSSEFAVIVEVHAAARSTLHPVGCEDDQSLSKYEFVVTSGSPVAADRVGP 480
TILNKIEAALTNQNLSVDVVDQCLVCLKEEWMNKVKVLFKFTKVDSRPKEDTQKLLSILG 540
ASEEDNVKLLKFWMTGLSKTYKSHLMSTVRSPTASESRN 579

UBL1/2 recognition motif: K/RxKxxxK
TRAF recognition motif: P/A/ExxS
```

**Figure S4** *FLCN contains multiple potential USP7 interaction motifs.*
The figure depicts the amino acid sequence of FLCN with the USP7 UBL1/2 (yellow) and TRAF recognition motifs (cyan) marked. The consensus sequences as defined by Kim and Sixma (Kim and Sixma, 2017) of the recognition motifs is given below. The x denotes any amino acid residue.

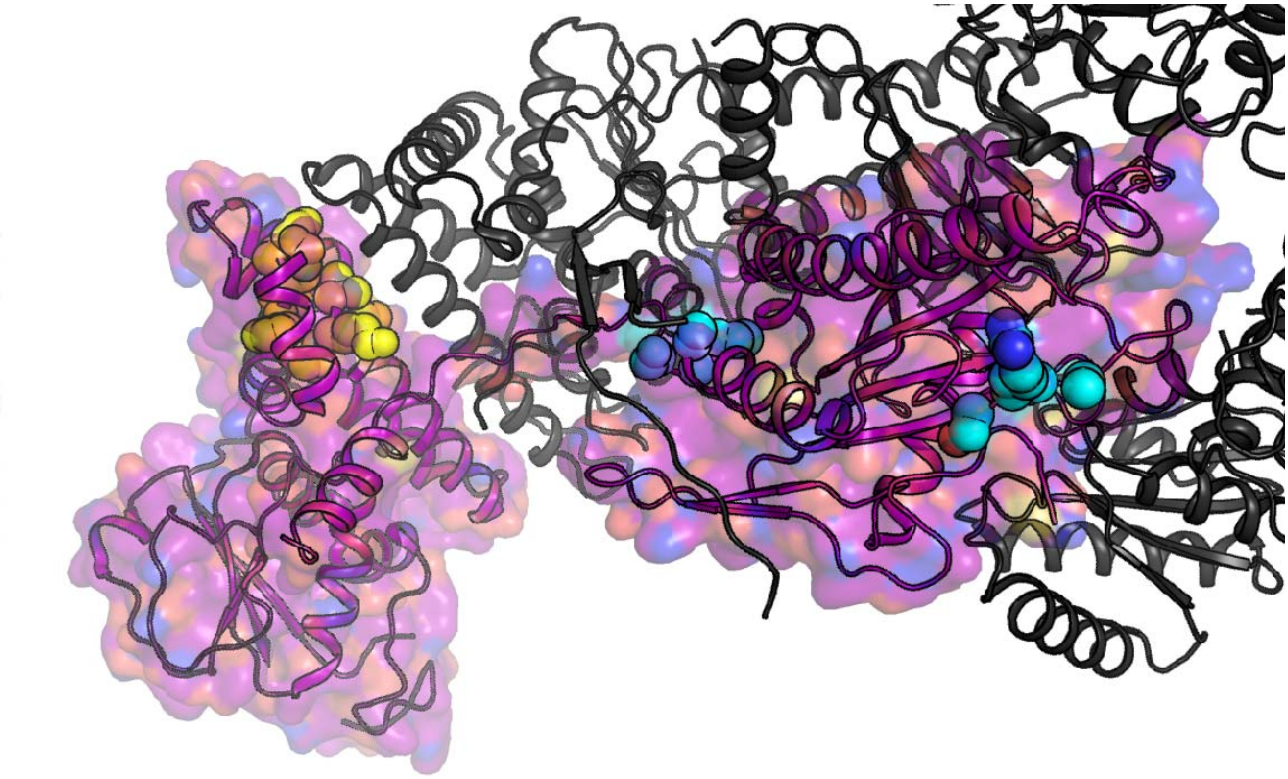

**Figure S5** *The potential USP7 binding sites in the FLCN structure.*

Mapping of the putative USP7 binding sites to the recently resolved cryo-EM structure of FLCN (PDB 6ULG) (Shen et al., 2019). Only 3 of the 10 putative sites correspond to regions that are resolved in the structure, namely 236AARS, 264ACGS, which are both in the N-terminal domain, and the UBL1/2 recognition motif (KVKVLFK) in the C-terminal domain. Two of these sites are covered by interaction interfaces with FNIP2 and are thus likely not accessible unless the complexes dissociates. Color coding as in Fig. S3.

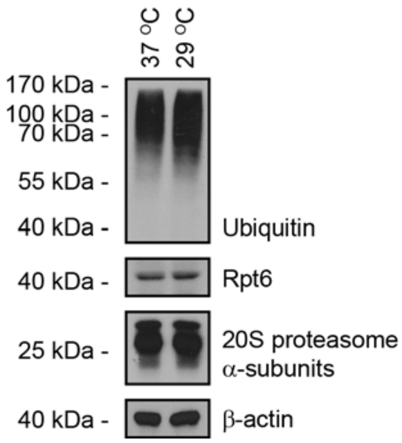


**Figure S6** *Normal appearance of the ubiquitin-proteasome system at 29 °C.*
The levels of ubiquitin-protein conjugates and 26S proteasomes was compared between U2OS cells grown at 29°C and at 37°C by blotting for ubiquitin, the 19S regulatory complex subunit Rpt6 and the 20S proteasome α-subunits. β-actin served as a loading control.
